# Imaging mitral/tufted glomeruli in the mouse olfactory bulb using the genetically encoded voltage indicator ArcLight

**DOI:** 10.64898/2026.09.22.753601

**Authors:** Lee Min Leong, Evan Lloyd, Douglas A. Storace

**Affiliations:** Department of Biological Science, Florida State University, Tallahassee, FL; Program in Neuroscience, Florida State University, Tallahassee, FL; Institute of Molecular Biophysics, Florida State University, Tallahassee, FL

**Keywords:** Olfactory bulb, mitral/tufted cells, glomeruli, 2-photon imaging, genetically encoded voltage indicator, ArcLight, fluorescent protein, mouse, optical imaging

## Abstract

In the mammalian olfactory bulb, each olfactory receptor neuron type maps to receptor-specific channels called glomeruli. These input axons interact with the apical dendrites of mitral/tufted cells, which are projection neurons that send axons to the olfactory cortex. Therefore, each glomerulus reflects the input-output relationship for a different olfactory receptor type, which is transformed by a complex synaptic network. While prior 2-photon Ca^2+^ imaging experiments have shown that the glomerular output is heterogeneous with respect to how odor concentration information is processed, the nature of voltage dynamics remains unclear. Therefore, we used the genetically encoded voltage indicator (GEVI) ArcLight to image the glomerular output *in vivo* in anesthetized mice. We found that ArcLight could resolve both excitatory and suppressive odor-evoked signals from mitral/tufted glomeruli. ArcLight expression remained stable over multi-week imaging sessions and successfully captured odor- and concentration-specific patterns of activation across a large concentration range. Importantly, this approach allowed us to resolve heterogeneous concentration-response sensitivities and respiratory-coupled dynamics. We also demonstrate that several newer GEVIs can similarly measure odor-evoked signals using epifluorescence imaging. Our study establishes ArcLight and other emerging GEVIs as powerful tools for dissecting how sensory information is encoded and transformed through the mouse olfactory bulb.

## Introduction

In the mammalian olfactory bulb (OB), odors are detected via the binding of volatiles to olfactory receptor neurons (ORNs), which drive varying degrees of activity across the receptor population. The ORNs project into receptor-specific channels called glomeruli, which map to different locations in the OB. Each glomerulus is innervated by the apical dendrites of mitral and tufted cells (MTCs), which send their axons to the olfactory cortex ^1, 2^. This input-output relationship is transformed by a complex synaptic network that includes different kinds of interneurons.

The ability to understand how sensory information is encoded and transformed by the OB network has been facilitated by the development of genetically encoded calcium and voltage indicators (GECIs and GEVIs, respectively). These protein-based optical reporters can be targeted to specific cell types and transduce changes in intracellular calcium or membrane potential into fluorescence changes ^3–5^.

Recent work performing 2-photon Ca^2+^ and glutamate imaging in MTC glomeruli revealed heterogeneous concentration-response relationships and temporal dynamics which are thought to facilitate the ability to distinguish between different kinds of odors ^4, 6–12^. However, the nature of voltage dynamics across the OB circuit remains unclear. We addressed this knowledge gap using the GEVI ArcLight to measure voltage dynamics from the glomerular output using *in vivo* 2-photon imaging, which minimizes out-of-focus signal. ArcLight is a strong candidate GEVI because it has been used to image different cell types within the mouse OB and can report odor-evoked activity using 2-photon imaging ^3, 5, 13^.

Here, we demonstrate that *in vivo* 2-photon imaging using ArcLight reliably resolves excitatory and suppressive odor-evoked voltage signals from individual mitral/tufted glomeruli. The ArcLight expression was bright and stable, which allowed for imaging from the same glomerular population across multi-week imaging sessions. Across the glomerular population, odor- and concentration-specific activation patterns could be detected even at relatively low odor concentrations. At the population level, MTC glomeruli exhibited a primarily monotonic concentration-response relationship. However, individual glomeruli exhibited heterogeneous responses, including those with monotonic or non-monotonic relationships with increasing odor concentration. Using standard resonant scanning speeds (∼31 Hz), ArcLight reliably reported respiratory-driven dynamics present in MTC glomeruli. Odors evoked significant increases in power at the respiratory frequency, although these changes were similarly heterogeneous across the glomerular population. Finally, we tested several newer-generation GEVIs under epifluorescence imaging conditions, finding that they are similarly capable of measuring odor-evoked activity from the mouse OB.

Overall, we find that the GEVI ArcLight has sufficient signal-to-noise to image biologically relevant dynamics even at near-threshold odor concentrations using 2-photon imaging. Our results establish ArcLight and other emerging GEVIs as high signal-to-noise tools capable of probing the temporal and spatial transformations that occur across the OB.

## Methods

### Ethical approval

All experiments were carried out according to the procedures and guidelines approved by the Florida State University Animal Care and Use committee under ethics approval reference #202100074.

### Animals

Both male and female adult C57BL/6, Tbx21-Cre (JAX stock #024507) and TH-Cre (JAX stock #008601) mice were used in this study. All transgenic mice used in the study (Tbx21-Cre and TH-Cre) were confirmed to express Cre recombinase via genotyping performed by Transnetyx (Cordova, TN).

### Surgical procedures

All mice included in this study were maintained in the FSU animal vivarium on a 12h/12h light/dark rhythm with *ad libitum* access to food and water. Male and female adult (>21 days) mice were anesthetized with ketamine/xylazine (90/10 mg/kg, IP, Zoetis, Kalamazoo, MI), placed on a heating pad and had ophthalmic ointment applied to their eyes. Anesthetic depth was monitored during all surgical procedures via pedal reflex. Mice were given a pre-operative dose of carprofen (20 mg/kg, SC, Zoetis, Kalamazoo, MI), atropine (0.2 mg/kg, IP, Covetrus, Dublin, OH), dexamethasone (4 mg/kg, IP, Bimeda, La Sueur, MN), and bupivacaine (1.5 mg/kg, SC, Hospira, Lake Forest, IL). Fur was removed from the top of the skull using a depilatory agent and rinsed, after which the skin was scrubbed with 70% isopropyl alcohol and iodine (Covidien, Mansfield, MA). An incision was made to remove the skin over the skull and blunt dissection was used to remove the underlying membrane. Dental cement (Metabond, Covetrus, Dublin, OH) was used to attach a custom headpost to the skull, which was held using a custom headpost holder. After the dental cement finished drying, a small craniotomy was made over the olfactory bulb (OB), into which a total of 500 nL of virus was injected as five 100 nL injections (Nanoject III, Drummond Scientific, Broomall, PA) distributed across five sites spaced evenly to promote uniform expression before sealing the cranial window with #1 cover glass.

Upon completion of the surgery, mice were allowed to recover on a heating pad until they were fully ambulatory. Animals were given a post-operative dose of carprofen (20 mg/kg, SC) at the end of the day of surgery and for at least 3 days post-operatively. The AAVs were allowed to express for at least 2 weeks before beginning imaging experiments. A subset of mice underwent a similar procedure except that a smaller craniotomy was made and after virus injection the incision was sutured. After identical post-operative treatment, similar surgical methods were used to install an optical window over the OB.

AAVs used in study were AAV1-CBA-ArcLight-nls-mCherry (3.7e11 vg/mL, Penn Vector Core, Philadelphia, PA), AAV1-CBA-ArcLightD-cre-ON-WPRE.SV40 (4.51e12 vg/mL, Penn Vector Core, Philadelphia, PA), AAV1-hSyn-DIO-ArcLightDco-WPRE (1.04e13 vg/mL, Obio Tech, Sugar Land, TX), AAV2/DJ-hSyn-JEDI-2P-Kv-WPRE-polyA (1.08e13 vg/mL, BrainVTA, Philadelphia, PA), AAV9-Syn-ASAP4e-kv (2.5e13 vg/mL), AAV9-EF1a-FLEX-ASAP4b(4g)-kv (1.8e13 vg/mL), AAV9-EF1a-FLEX-ASAP3 (2.0e13 vg/mL).

### Histology

Mice were euthanized via an IP injection of Euthasol and either underwent cardiac perfusion with phosphate-buffered saline and 4% paraformaldehyde or had their brains extracted and post-fixed in 4% paraformaldehyde before being sectioned on a vibratome in 40 µm sections (Leica VT1000S, Deer Park, IL). OB sections were mounted on slides and coverslipped using Fluoromount-G containing DAPI (SouthernBiotech, Birmingham, AL). Endogenous fluorescence expression of the GEVIs was observed using a GFP filter set on a Zeiss Axioskop epifluorescence microscope and was imaged on a Nikon CSU-W1 spinning disk confocal microscope using a 20x 0.95 N.A. objective lens.

### Olfactometry

For all 2-photon imaging experiments, the odorants methyl valerate (99% pure, CAS #624-24-8, Sigma-Aldrich, #148997), isoamyl acetate (99% pure, CAS #123-92-2, Thermo Scientific #150662500), acetophenone (99% pure, CAS #100-52-7, Thermo Fisher Scientific #AAA12727AP), and 2-heptanone (99%, CAS #110-43-0, Thermo Fisher Scientific #A10200AE) were diluted in mineral oil to concentrations of 0.0077%, 0.0463%, 0.278%, 1.667%, 10%, and 60%. The liquid diluted odors were delivered at 10% of saturated vapor using an olfactometer in which an air pump constantly delivered clean air to two mass flow controllers (MC-100SCCM, and MC-1SLPM, Alicat, Tucson, AZ). The MC-100SCCM controlled air flow through the odor vials connected in line through a pair of 4-valve manifolds (360T081, NResearch, West Caldwell, NJ) at 50 mL/min, which blended with clean air flow at 450 mL/min controlled by the MC-1SLPM. The resulting odorized air stream connected to a dual 3-way solenoid valve (360T041, NResearch, West Caldwell, NJ) which was connected to an exhaust, a flow-rate matched clean air stream and a Teflon delivery manifold which served as the final delivery apparatus placed in front of the mouse’s nose. The 3-way solenoid valve sent the odorized air stream to the exhaust and the clean air stream to the mouse prior to the odor trigger. Triggering the 3-way solenoid caused the odor to be injected into the delivery manifold. The odor delivery time-course for both olfactometer setups was confirmed using a photoionization detector (200C, Aurora Scientific, Aurora, ON).

### Imaging procedures

Prior to data collection mice were positioned underneath the microscope and the headpost holder angle was adjusted to optimize the imaging field of view. During data collection, head-fixed mice were placed underneath the microscope objective with the olfactometer placed in front of the animal’s nose. A pressure sensor embedded in the olfactometer measured respiration signals which were amplified and low-pass filtered using a differential amplifier (Model 3000, AM-Systems, Sequim, WA) and recorded by the imaging system.

For 2-photon imaging experiments (**Figures 1-6**), odors were delivered in 6 concentration steps between 0.0077% and 60% of pure odor diluted in mineral oil at 10% of saturated vapor, alongside air trials in which the odor stream was passed through a previously unused vial. For epifluorescence imaging (**Figure 7**), odors were delivered at concentrations between 6 – 10% of saturated vapor of undiluted odor. The exception was JEDI-2P, for which 10% of pure odor was diluted in mineral oil and delivered at 10% of saturated vapor. Each odor-concentration pairing was measured in response to a minimum of 3 single trials separated by 3 minutes. For each animal, we prioritized measuring multiple repetitions for each concentration for a particular odor before a second odor was attempted.

**Figure 1:**
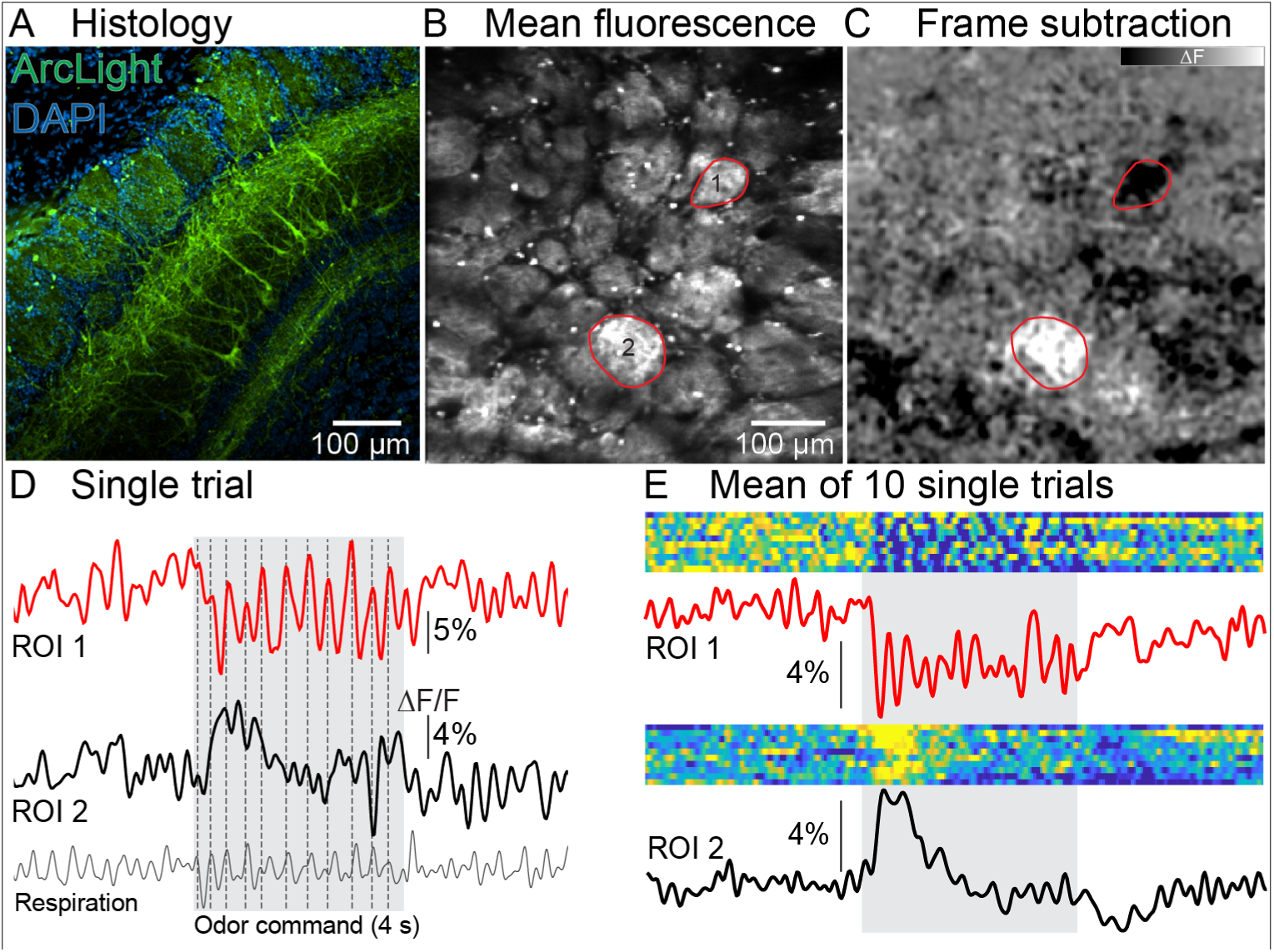
*In vivo* 2-photon imaging from ArcLight targeted to mitral/tufted glomeruli. (A) Histology from a Cre-dependent AAV injection into a Tbx21-Cre transgenic mouse. (B-C) Mean fluorescence (B) and a frame subtraction (C) illustrating the odor response pattern (right) from an exemplar preparation. (D-E) Single trial (D) and mean (E) fluorescence time course from 2 glomeruli in response to odor stimulation.

2-photon imaging was performed using a Sutter MOM 2-photon microscope equipped with an 8 kHz (30.9 Hz) resonant scanner (Cambridge Technology, USA) and an emission pathway equipped with a GaAsP PMT (#H10770PA-40-04, Hamamatsu, Japan). Laser excitation was provided using an Alcor 920-2W laser with an internal power modulator. Imaging was performed using a 10x 0.5 N.A. objective lens (Yu et al., 2024). Laser power was confirmed to be less than 150 mW at the output of the objective lens measured using a power meter (Newport 843-R) for a scanning area of 1138 μm^2^.

Epifluorescence imaging was carried out using a custom microscope built from Thorlabs (Newton, NJ) components with a 35 mm CCTV lens, or a 10x 0.45 N.A. objective. Illumination was provided by a Prizmatix LED (UHP-T-LED-455, Holon, Israel) filtered with a 488/10 nm filter (FF01-488/10, Semrock, West Henrietta, NY). A dichroic mirror (59009bs, Chroma, Bellows Falls, VT) directed the excitation light toward the preparation and transmitted the fluorescence emission through a band-pass filter (59009m, Chroma, Bellows Falls, VT), before being recorded using a DaVinci 1k camera (RedShirtImaging, Decatur, GA). Imaging was carried out between 40 and 100 Hz at a spatial resolution of 256x256 pixels.

### Data analysis

#### Frame Subtraction Analysis

The mean fluorescence images are generated from the average of all the frames during the imaging trial, or all the frames prior to odor stimulation (**Figures 1B**, **2A, S1A**). The frame subtraction images were generated by subtracting the 1-2 seconds during odor stimulation from the pre-odor baseline (**Figures 1C**, **2A, S1B**). The frame subtraction intensity scales were adjusted to optimally display peaks of activation.

**Figure 2:**
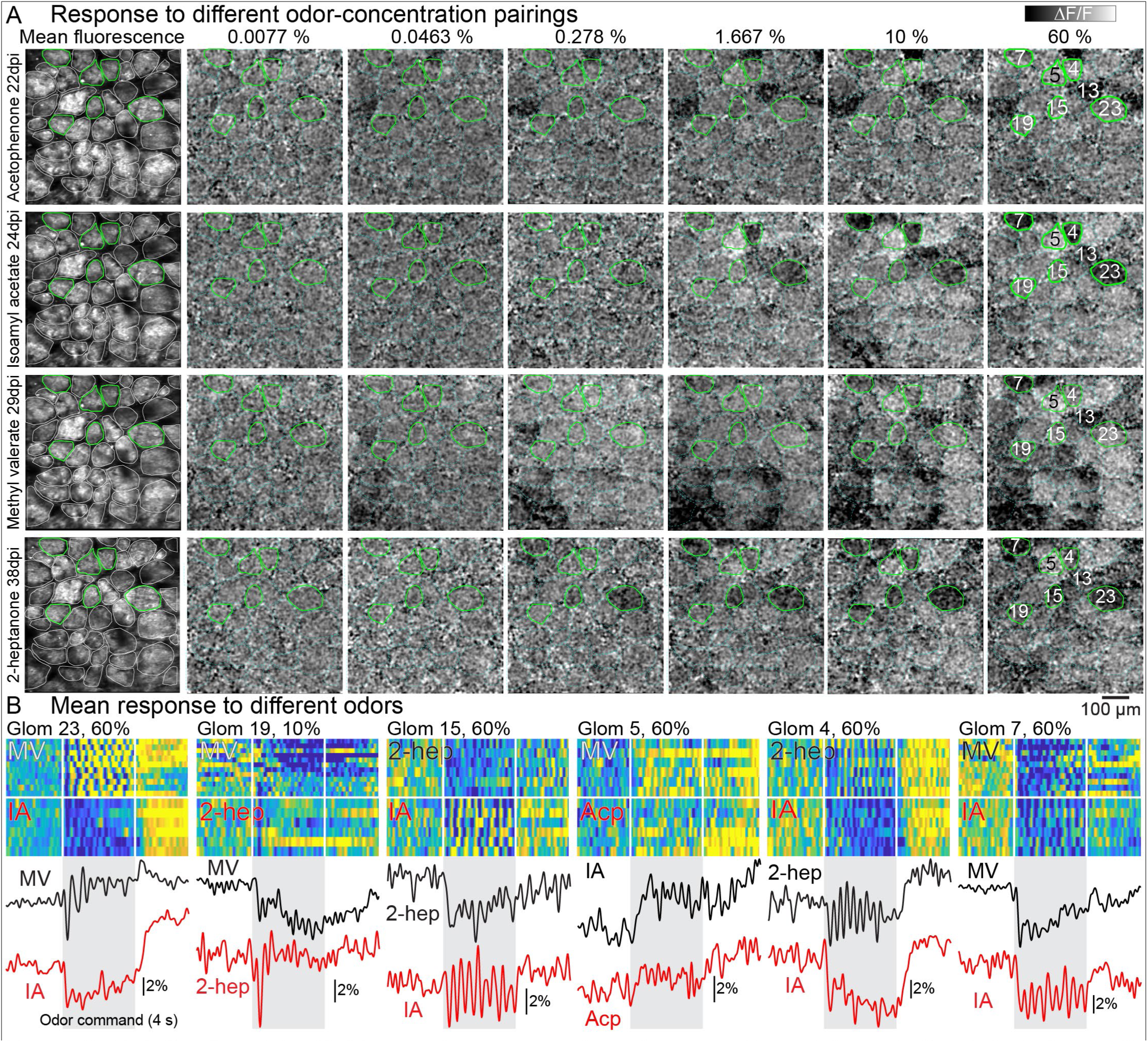
(A, left column) ArcLight mean fluorescence from the same field of view of mitral/tufted glomeruli imaged between 22-38 days post injection (dpi). (A, right columns). Frame subtractions images illustrating the response to four odors (rows) at 6 concentrations (columns). (B) The response from 6 glomeruli to different odor pairs. MV, methyl valerate; IA, isoamyl acetate; Acp, acetophenone; 2-heptanone, 2-hep.

#### Processing and segmentation

For 2-photon imaging data, individual glomeruli were segmented based on morphological characteristics in the mean fluorescence and peaks of activity in the frame subtraction analysis (**Figure 1B-C**, *red polygons*) in Turbo-SM (SciMeasure, Decatur, GA). For epifluorescence imaging data, glomeruli were defined based only on the presence of glomerular sized peaks of activity in a frame subtraction analysis. The pixel areas containing the regions of interest were saved and the fluorescence time course values from each region of interest were extracted for subsequent analysis. For all traces, fluorescence was converted to ΔF/F by dividing the raw fluorescence values by the mean of all the frames prior to the odor command.

#### Other analyses

Individual trials were aligned to the first inhalation following the odor command trigger and were low-pass filtered between 4-8 Hz. Averaged traces and corresponding measurements for each odor-concentration condition were performed in inhalation-aligned averaged trials. Odor response amplitudes (ΔF/F) were calculated by measuring the largest difference between a moving window average of 192 msec during the first 500 msec following inhalation of the odor, and the 2 seconds prior to odor stimulation. This peak change was selected for glomeruli in which the change exceeded 3 standard deviations from the baseline (**Figure 3A**). To avoid the selection of spurious peaks for non-responsive glomeruli, the amplitude for glomeruli with changes less than 3 standard deviations were calculated as the mean of the entire 500 msec period following the inhalation of the odor. Glomeruli were subsequently defined as responsive if they exhibited an odor response with a change greater than 3 standard deviations at the highest tested concentration (60% liquid dilution), and response amplitudes were quantified at all concentrations in these glomeruli. Rise time was defined as the time from 10% to 90% of the peak response amplitude following odor inhalation (**Figure 3A-B**). Response thresholds are defined as the lowest concentration that evoked an odor-evoked change that was more than 3 standard deviations above baseline (**Figure 3E**). The error bars in the concentration-response relationships in **Figure 4** illustrate the standard error of the mean of the single trial measurements for that odor-concentration condition.

**Figure 3:**
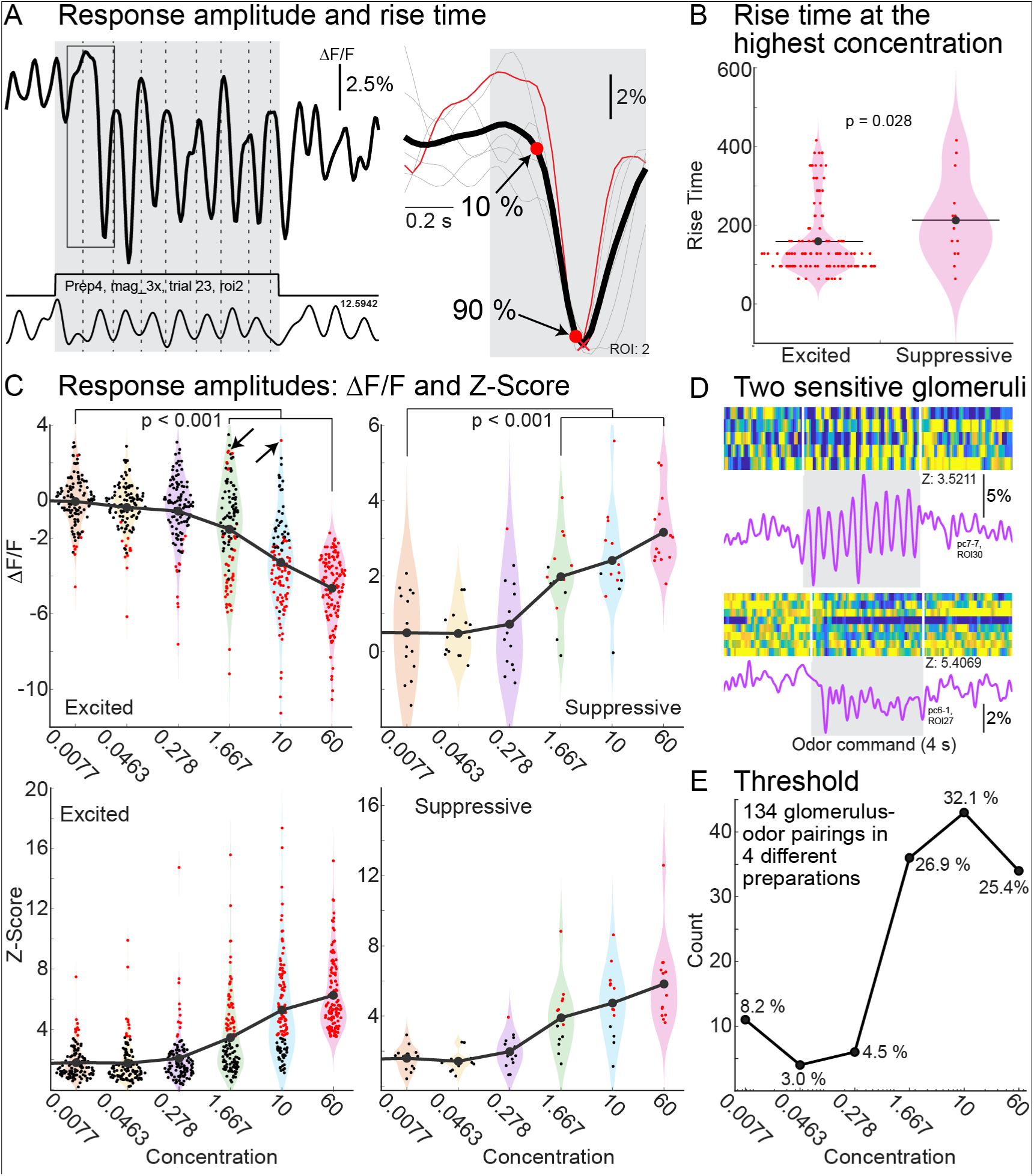
(A) Single trial example of excited response (left) and a time expansion of a 6 single trial sniff aligned average (right). The X and the circle indicate the peak odor response and the 10 to 90% rise time. (B) Rise time of odor responses for all glomeruli at 60% odor dilution. (C-D) Mean response amplitudes in ΔF/F (top) and Z-Score (bottom) for each concentration. Red markers indicate glomeruli with Z-Scores greater than 3.5. The arrows at 1.667 % and 10 % indicate excitatory glomeruli with suppressive responses at an intermediate concentration. (D) Single trial (heat maps) and mean fluorescence time course measurements from 2 glomerulus-odor pairings at the lowest tested concentration (0.0077%). (E) The threshold concentration for all glomerulus-odor pairings.

**Figure 4:**
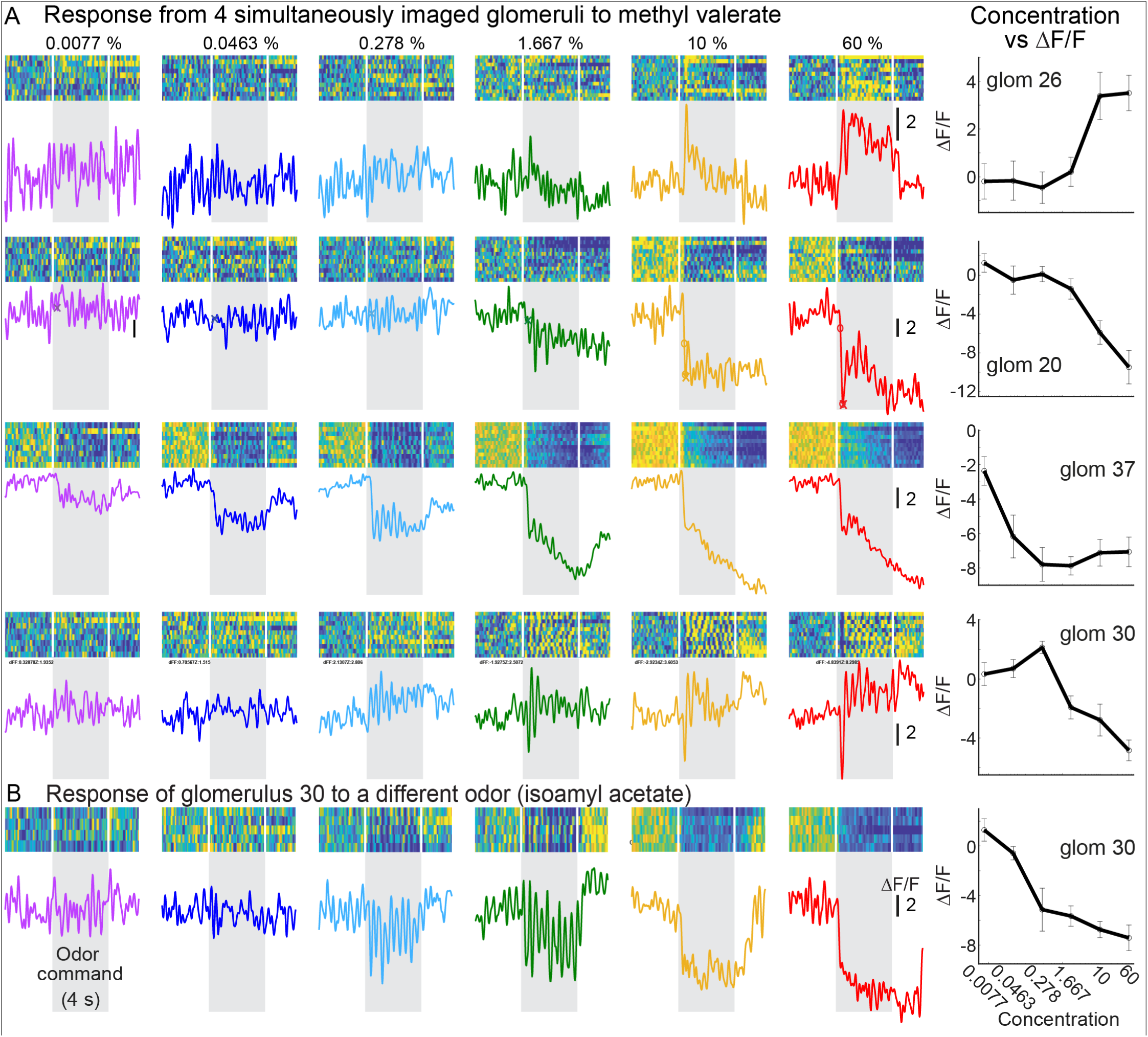
Monotonic and non-monotonic concentration-response relationships in mitral/tufted glomeruli. (A) Four different glomeruli simultaneously imaged in the same field of view to a range of concentrations of the same odor. (left columns) Single trial and mean fluorescence time course at different concentrations. (right-most column) Mean concentration-response relationship for each glomerulus. The errorbars indicate s.e.m. (B) Response from glomerulus 30 (4th from top) to a different odor.

#### Respiration quantification

The population FFT in **Figure 5B** was generated by averaging the FFT of each individual single trial for each concentration, which were subsequently averaged together. Similar results were obtained by averaging the respiration signal across all single trials. The shuffling analysis involved randomly adjusting the alignment of the respiration aligned trials between 0-352 msec. This was repeated 500 times, and the resulting traces were averaged together (**Figure 5D-F**). A Fast Fourier Transform (FFT) was performed on the respiration measurements (**Figure 5B-C**) and the ArcLight signal (**Figure 5D-F**, **Figure 6**) using the fft function in MATLAB ^13^. Power at the respiration frequency was integrated between 2-4 Hz based on a visual analysis of the respiration traces and the peak of the respiration FFT (**Figure 5A-B**).

**Figure 5:**
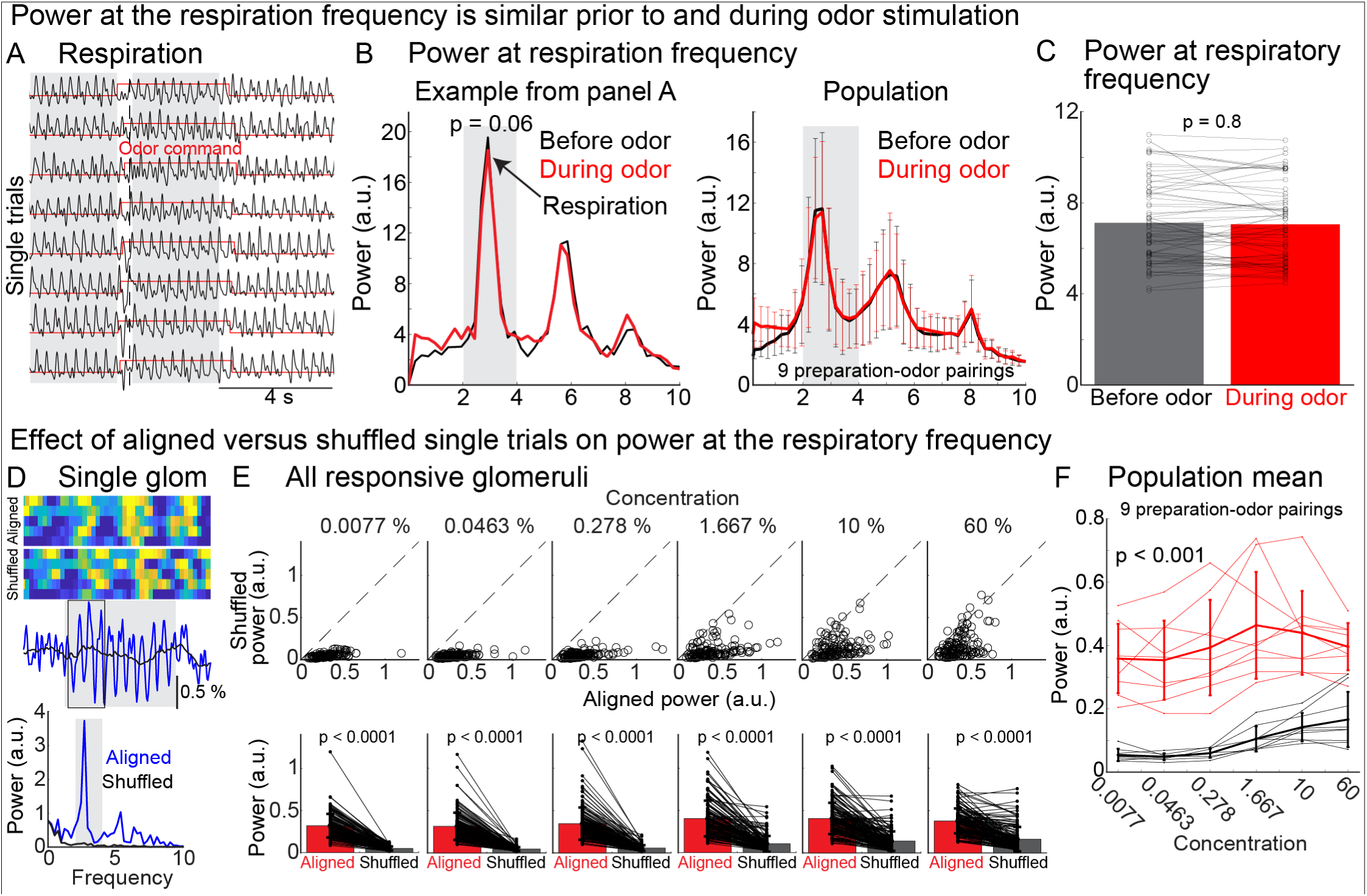
(A) Exemplar respiration traces during imaging trials. (B) Power at the respiration frequency for the example in panel A (left) and for the population (right). (C) The mean power at the respiration frequency before and after odor presentation. The vertical gray bars in panels A-B illustrate the before and after time points used for power calculations (A), and the integrated frequency range (B). (D) Single trial (top), mean fluorescence (middle), and the power at the respiration frequency (bottom) for one glomerulus using respiration aligned trials (colored) versus shuffled alignment (black). (E) Power at the respiration frequency using respiration aligned versus shuffled alignment for all glomeruli. (F) Mean power at the respiration frequency for each preparation before and after shuffling respiration alignment.

**Figure 6:**
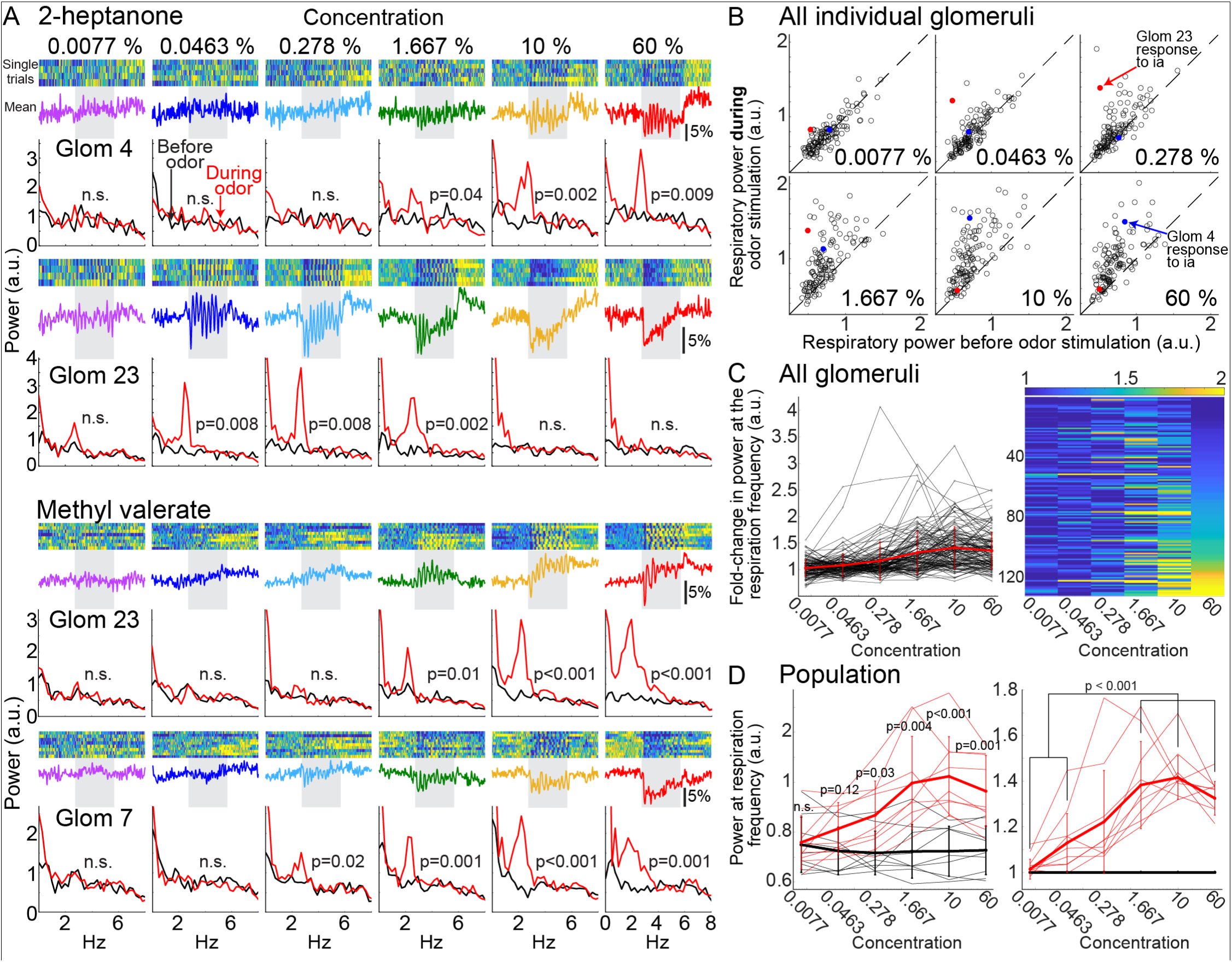
(A) Single trial (top), mean fluorescence time course (middle) and frequency spectra (bottom) before and after the odor presentation from two glomerulus-odor pairings simultaneously imaged from the same field of view. Both pairs are from the same mouse preparation. (B) Power at the respiration frequency before and after odor presentation for all responsive glomeruli at each concentration. The exemplar glomerular responses to 2-heptanone are illustrated as blue and red circles (glom 4 and 23, respectively). (C) Fold-change in power at the respiration frequency before versus after the odor presentation for each glomerulus plotted as a line graph and heat map. (D) Mean power (left) and fold-change in power (right) at the respiration frequency before (black) and after (red) odor presentation for each preparation-odor pairing.

#### Epifluorescence recordings

The fluorescence time course traces were low-pass filtered and had an exponential drift subtracted using MATLAB. The fluorescence brightness analysis was measured by choosing a circular region of interest over the brightest area of the right OB. Regions of interest diameters ranged from 100 to 180 µm approximately, depending on whether the recording was made with the 35 mm CCTV lens or the 10x 0.45 N.A. objective. The response amplitude was measured using the peak-to-peak value of the largest change following the first inhalation of the odor from the mean odor-evoked fluorescence response trace plotted in MATLAB for a caudal and a rostral region of interest. The signal-to-noise ratio was calculated by dividing the response amplitude values by the peak-to-peak value of the baseline noise in the 500 msec prior to the odor trigger. For each GEVI, preparations that yielded no detectable odor-evoked signal (recorded as zero values) were excluded from descriptive statistics and formal statistical comparisons to accurately assess the functional metrics of responding cells. However, these zero values were retained in all visual representations to provide an overview of the preparations.

#### Statistical Analyses

All statistical comparisons involved non-parametric tests including the Wilcoxon rank-sum test, and the Kruskal-Wallis test (ranksum and kruskalwallis functions in MATLAB). A ranksum test was used for statistical differences between pairs of GEVI constructs with a Bonferroni correction applied for multiple group comparisons. Specifically, for the group of constitutively expressed GEVIs (ArcLight, ASAP4e-kv, and JEDI-2P-kv), the threshold for significance was adjusted to p < 0.0167 (0.05 / 3).

## Results

### ArcLight reports odor-evoked activity in mitral/tufted (MTC) glomeruli using 2-photon imaging

To test whether ArcLight could be used to measure odor-evoked signals from MTC glomeruli using *in vivo* 2-photon imaging, we injected a Cre-dependent adeno-associated virus (AAV) into Tbx21-Cre transgenic mice. In histological sections, these preparations exhibited expression in MTCs and their apical dendrites in glomeruli (**Figure 1A**). During *in vivo* 2-photon imaging in anesthetized mice, individual glomeruli were detectable based on their mean fluorescence and localized odor-evoked fluorescence changes (**Figure 1B-C***, red polygons*). Example single trials from two glomeruli, along with their corresponding inhalation-aligned mean fluorescence time courses, illustrate that odors evoked robust, stimulus-locked, and sometimes respiratory-coupled fluorescence changes (**Figures 1D-E**). Odor responses included both fluorescence decreases and increases, which reflect depolarization and hyperpolarization, respectively, that were consistent across individual trials separated by 3-minute inter-trial intervals (**Figure 1D-E**, red vs black trace). Qualitatively similar odor responses were evident in 9 of the 10 anesthetized preparations. To confirm the utility of ArcLight across different physiological states, we also performed imaging in awake, head-fixed mice. Despite the expected state-dependent movement artifacts, odor-evoked signals remained detectable in 4 of 5 preparations (**Figure S1**). Therefore, ArcLight is suitable for measuring *in vivo* odor-evoked voltage signals from MTC glomeruli in the mouse olfactory bulb using 2-photon imaging.

### ArcLight fluorescence and odor-evoked signals remain stable across different imaging sessions

ArcLight fluorescence in MTC glomeruli was stable over time, which allowed for the reliable re-alignment of the same glomeruli across consecutive imaging sessions (**Figure 2A**, representative mean fluorescence across days 22, 24, 29 and 38 days *post injection*). We imaged 7 of the 10 previously described preparations on multiple days, yielding a total of 35 imaging sessions (5 ± 2.77 sessions per preparation, range of 2 to 9; mean ± standard deviation) covering a span of 15 ± 13.4 days (range of 2 to 35 days). Importantly, odor-evoked signals could still be measured during each imaging session across all 7 preparations. Therefore, virally driven ArcLight enables longitudinal voltage imaging *in vivo*.

### ArcLight can detect odor- and concentration-specific patterns of activation in MTC glomeruli

We measured responses to different odors at concentrations between 0.0077% and 60% (percent liquid dilution in mineral oil) in 4 of the 9 preparations with responsive glomeruli (**Figure 2**). Odor-evoked activation maps revealed that the number of responsive glomeruli increased with increasing odor concentration within the same field of view (**Figure 2A**, compare patterns within a row). Similarly, different odors evoked different patterns of activation across the glomerular ensemble (**Figure 2A**, compare patterns within each column).

Exemplar odor responses from 6 different glomeruli to different odors illustrate the typical signal-to-noise ratio measured in these experiments, and that different odors could also elicit distinct time course dynamics and varying degrees of respiratory-coupling within the same glomeruli (**Figure 2B**, compare the response of Glom 15 and Glom 4 to 2-heptanone and isoamyl acetate, respectively). Therefore, the signal-to-noise ratio of ArcLight allows for resolving the spatial and temporal dynamics of MTC glomeruli across a range of concentrations.

We imaged 152 individual glomeruli across the 4 mouse preparations (38 ± 6.7 glomeruli per preparation; range of 29-44). A subset of these were imaged in response to multiple odors (methyl valerate, isoamyl acetate, acetophenone and 2-heptanone), which yielded 351 unique glomerulus-odor pairings. Each odor pairing was sampled across all 6 tested concentrations plus a clean air control, yielding a total of 2457 glomerulus-stimulus pairings.

We quantified the response rise time (the time from 10% to 90 % of the peak response) and the mean concentration-response relationship using the largest change following the first inhalation of the odor (**Figure 3A**). We defined responsive glomeruli as those exhibiting a minimum of a 3 standard deviation change from the pre-odor baseline at the highest tested concentration. At the highest tested concentration (60%), excited glomeruli exhibited significantly faster rise times than suppressive glomeruli (excited: 161 ± 92 msec; suppressed: 212 ± 107 msec, ranksum test p = 0.028) (**Figure 3B**). At the population level, increasing the odor concentration evoked monotonic increases in response amplitude (ΔF/F). The three highest tested concentrations (1.667%, 10%, and 60%) had significantly larger signals in comparison to the lowest tested concentration (0.0077%) (**Figure 3C**, Kruskal-Wallis test with Tukey post-hoc: excited: p< 0.0001; suppressive: p<0.0001; red data points indicate an odor response with a 3 standard deviation change at that concentration).

Measurements from 2 representative glomeruli at 0.0077% illustrate that ArcLight can report odor-evoked activity even at relatively low concentrations (**Figure 3D**). We calculated the threshold concentration for each glomerulus, defined as the lowest concentration evoking a minimum of a 3 standard deviation change above baseline (**Figure 3E**). Across all responsive glomeruli, ∼11% responded at these relatively low concentrations, with most beginning to respond at 1.667% and 10% (**Figure 3E**).

While the population response amplitude scaled monotonically, individual glomeruli imaged simultaneously within the same field of view displayed notable heterogeneity in how they responded to concentration changes. Some glomeruli exhibited monotonically increasing relationships where increasing concentrations evoked progressively larger excitatory or suppressive response amplitudes (**Figure 4A**, glomeruli 26 and 20). Other glomeruli exhibited non-monotonic concentration-response relationships which included those that responded most strongly to an intermediate concentration, with further increases in concentration evoking progressively weaker responses (**Figure 4A**, glomerulus 37). Other non-monotonic glomeruli responded with a suppressive response at some concentration but transitioned to an excitatory response at the highest concentration (**Figure 4A**, Glom 30 response to methyl valerate; also see **Figure 3C**, the black arrows at 1.667% and 10% point to two glomeruli with this property). The concentration-response relationship of the same glomerulus to 2 different odors illustrate that the manner in which a glomerulus responds to concentration changes is odor-specific (**Figure 4A-B**, Glom 30). Therefore, ArcLight can resolve heterogeneous and odor-specific concentration-response dynamics across individual MTC glomeruli within the same field of view.

### ArcLight reliably reports respiratory-coupled signals using 2-photon imaging

Exemplar respiration measurements and the corresponding spectral analysis from an individual preparation demonstrate that our ketamine/xylazine anesthetized mice exhibited relatively slow and stable respiratory rates between 2-4 Hz before and during the odor (**Figure 5A**). Across the population, the power within the respiratory frequency range (2-4 Hz) did not significantly differ before and during the odor presentation (**Figure 5B-C**; ranksum test; p = 0.8).

When single trials were aligned to the first inhalation during the odor presentation, the ArcLight signal in many glomeruli tracked the respiration frequency and produced corresponding peaks in spectral power (**Figure 5D**, *blue trace*). Randomly shuffling the single trial alignment significantly reduced the phasic oscillations and the power at the respiratory frequency in these examples (**Figure 5D**, *blue versus black traces*). Shuffling similarly reduced power at the respiration frequency at all concentrations at the individual glomerular level and when averaged across preparations (**Figure 5E-F**). Therefore, ArcLight reliably reports respiratory-coupled voltage signals in MTC glomeruli.

Individual glomeruli imaged within the same field of view exhibited complex relationships between concentration and respiratory coupling. Some glomeruli had a monotonic relationship between increases in concentration and power at the respiratory frequency (**Figure 6A**, response of glomeruli 4 and glom 23 to 2-heptanone and methyl valerate, respectively). Other glomeruli exhibited strong respiratory coupling at an intermediate concentration but adapted and became uncoupled at higher concentrations (**Figure 6A**, response of glomeruli 23 and 7 to 2-heptanone and methyl valerate, respectively). The same glomerulus could exhibit respiratory-coupled or adapting responses in an odor-specific manner (**Figure 6A**, compare the response of glomerulus 23 to 2-heptanone versus methyl valerate).

The relationship between concentration and power at the respiratory frequency is illustrated for all responsive glomeruli in the data set (**Figure 6B-C**). Across the population, odors evoked significant increases in respiratory power relative to baseline levels at the four highest concentrations (**Figure 6D**, left). While the normalized change in power trended downward at the highest concentration, the mean change was only significantly different when compared to the lowest tested concentrations (**Figure 6D**, right).

### Other GEVIs report odor-evoked activity in the mouse olfactory bulb using epifluorescence imaging

We tested the ability of several newer generation GEVIs to report odor-evoked signals from the mouse OB in anesthetized mice using epifluorescence imaging. The tested sensors belong to two structurally distinct families, with the ArcLight family fusing the voltage-sensing domain of *Ciona intestinalis* voltage-sensing phosphatase to super ecliptic pHluorin carrying a A227D mutation, and the ASAP/JEDI family inserting a circularly permutated GFP into the extracellular S3-S4 loop of the chicken (*Gallus gallus*) voltage-sensing domain (Evans et al., 2023, Liu et al., 2022, Villette et al., 2019).

ArcLight, ASAP3, ASAP4b, ASAP4e and JEDI-2P were either expressed constitutively or in a Cre-dependent manner in TH-Cre or Tbx21-Cre transgenic mice via adeno-associated virus injections into the OB. In preparations with constitutive expression, odor-evoked signals were detectable in 4 of 5 preparations for ArcLight, 5 of 7 preparations for ASAP4e-kv, and 2 of 3 preparations for JEDI-2P-kv (**Figure 7A**). For Cre-dependent injections into TH-Cre mice, odor-evoked signals were detectable in 6 of 7 ArcLight preparations and 3 of 3 ASAP3 preparations (**Figure 7B**). In Tbx21-Cre mice, signals were detected in 5 of 6 ArcLight preparations, and 1 of 5 ASAP4b-kv preparations (**Figure 7C**).

**Figure 7:**
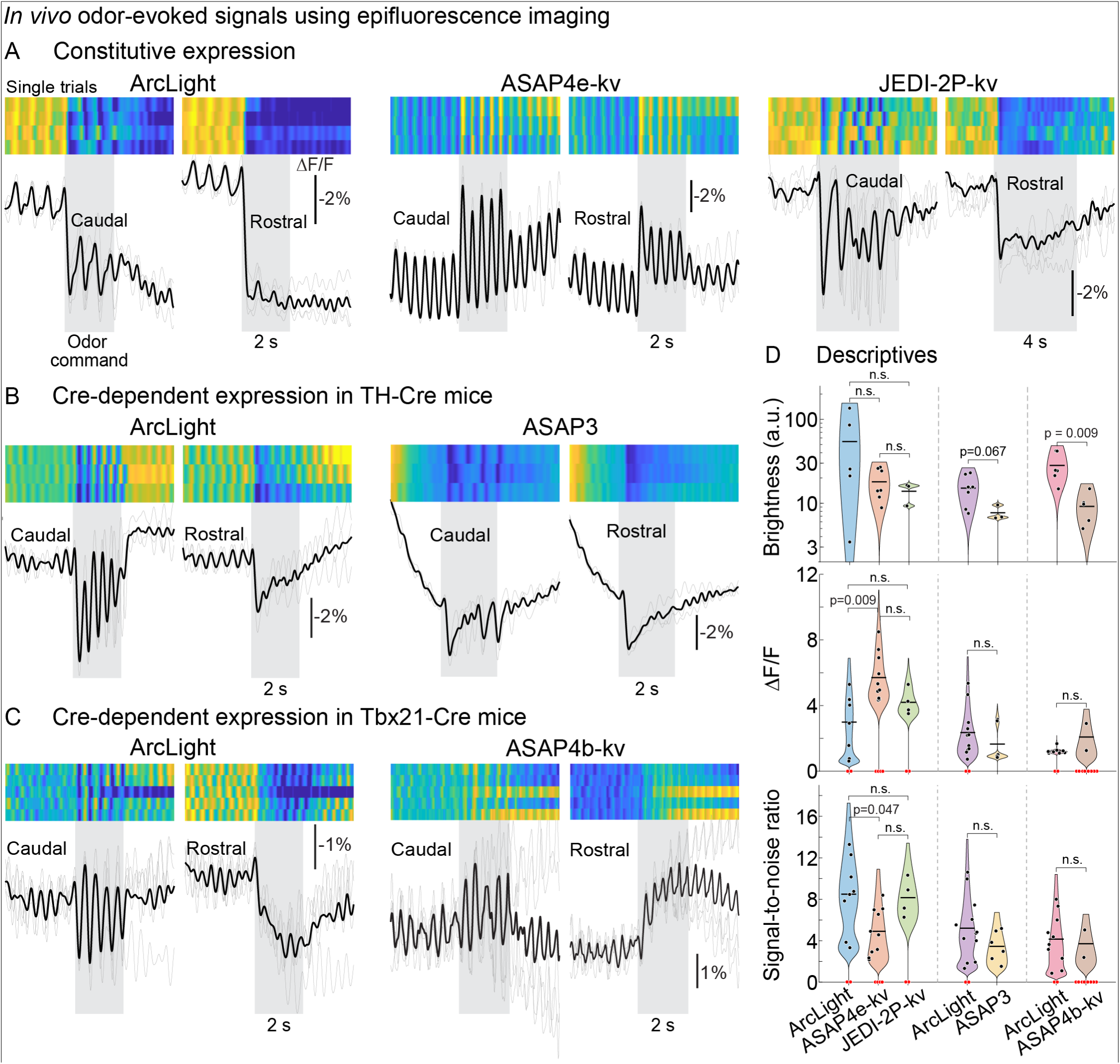
*In vivo* odor responses in the mouse olfactory bulb from different GEVIs imaged using epifluorescence. (A-C) Virally-driven constitutive (A) and Cre-dependent (B-C) expression in two different transgenic lines (B-C). The single trial (top) and mean fluorescence (bottom) is illustrated for each tested GEVI from a caudal (left) and rostral (right) region of interest. (D) Mean brightness (top), response amplitude (middle), and signal-to-noise ratios (bottom) for each GEVI. Black points indicate measurements from responsive preparations. Tested preparations that did not exhibit an odor response are indicated in red and are excluded from the statistical comparisons. The dashed lines separate constructs expressed constitutively (left), in TH-Cre mice (center) and in Tbx21-Cre mice (right). P-values are from two-sided Wilcoxon rank-sum tests, Bonferroni-corrected within the constitutively expressed group.

Notably, each sensor was capable of detecting the temporal heterogeneity present in epifluorescence imaging across the rostral-caudal axis of the bulb, in which glomeruli located caudally demonstrate greater respiration-locking than those located rostrally (**Figure 7A-C**, compare both ROIs for each GEVI comparison) ^13–15^.

ArcLight was brighter on average than the other GEVIs, with a small but statistically significant difference in comparison with ASAP4b-kv in Tbx21 mice (**Figure 7D**, top). In constitutive experiments, ASAP4e-kv exhibited the largest odor-evoked response amplitudes (ΔF/F). However, its larger baseline peak-to-peak resulted in a similar signal-to-noise ratio to ArcLight and JEDI-2P (**Figure 7D**, middle and bottom panels). Overall, these results demonstrate that multiple GEVIs provide sufficient signal-to-noise ratio to detect odor-evoked signals from the mouse OB *in vivo* using epifluorescence imaging.

## Discussion

Here we demonstrate that the GEVI ArcLight has sufficient signal-to-noise ratio to measure respiratory-coupled, odor-evoked signals from mitral/tufted (MTC) glomeruli in the mouse olfactory bulb when using *in vivo* 2-photon imaging. Odor-specific excitatory and suppressive responses were measured in different MTC glomeruli and could be reliably imaged across multi-week sessions. This is important because it enables longitudinal experiments that can track how sensory representations evolve over time, or in response to different experimental conditions (e.g., learning or metabolic state). Additionally, the improved spatial resolution of 2-photon imaging allowed us to resolve functionally heterogeneous response characteristics present across glomeruli in the same imaging field of view that were previously detected using calcium and glutamate imaging. Finally, we used epifluorescence imaging to demonstrate that several newer-generation GEVIs are similarly capable of reporting odor-evoked activity from different cell populations in the mouse olfactory bulb. Our results highlight the potential for GEVIs to become a powerful tool for understanding how sensory information is encoded and transformed within the mouse olfactory bulb.

### Comparison with previous studies

Here we extended our previous work using ArcLight by performing 2-photon imaging from MTC glomeruli and finding the presence of suppressive odor responses and heterogeneous concentration-response relationships (**Figures 1**-**4**). While both phenomena have been reported using calcium and glutamate sensors ^8, 9, 11, 16^, they were not previously reported in datasets collected using epifluorescence imaging ^4, 6^. We propose that the use of 2-photon imaging facilitated our ability to detect these changes by reducing the amount of out-of-focus fluorescence that can obscure responses from individual glomeruli.

The finding that ArcLight can resolve respiratory-coupled dynamics from MTC glomeruli is consistent with reports that mitral cell spiking activity is linked to the respiratory cycle ^17–19^. These dynamics were also reported in prior measurements with ArcLight performed using epifluorescence imaging, and 2-photon imaging line scans using a slower, non-resonant scanner ^13^. Therefore, our data extend these results to demonstrate that 2-photon microscopy with a resonant scanner has sufficient temporal resolution to detect respiratory-coupled oscillations even when imaging a relatively large field of view that includes many glomeruli (**Figures 5**-**6**) ^20^.

Our observation that the power at the respiration frequency is not only heterogeneous across glomeruli but also increases monotonically at the population level demonstrates that respiratory coupling has a complex relationship with odor concentration. While this result aligned with previous studies demonstrating that different MTC glomeruli exhibit distinct time course dynamics ^13–15^, glutamate release onto MTC glomeruli can be reduced at high concentrations for some odors ^11^. While the power at the respiration frequency trended downward at the highest concentration (**Figure 6**), the effect was not significant primarily because increasing the odor concentration recruited many strongly respiratory-coupled glomeruli that were inactive at lower concentrations (e.g., glomerulus 4 in **Figure 6A**). Future studies will need to address whether respiratory coupling is driven primarily by the sensitivity of the ORN input itself, or whether it reflects processing within the bulb ^8^.

Finally, the previous studies that imaged the olfactory bulb using ArcLight used high odor concentrations that exceed those found in many natural sources ^6, 13, 21^. Therefore, our ability to reliably measure odor responses even at relatively low concentrations (0.0077%) demonstrates that ArcLight has the sensitivity to detect odor-evoked activity at odor concentrations approaching the thresholds of olfactory perception in mice for a subset of the tested odors (**Figure 3C-D**) ^22–24^.

### Methodological considerations

Here we find that at the highest tested concentration (60%), MTC glomeruli with suppressive odor responses have slower rise times than those with excitatory responses (**Figure 3B**). Future studies are needed to confirm this result with a larger pool of suppressive responses, and to determine whether it applies broadly or if it occurs in a concentration-dependent manner. However, this observation raises the possibility that ArcLight, and GEVIs in general could be useful for resolving responses that may be driven by olfactory bulb circuitry. While most of our experiments were performed in anesthetized mice, we confirmed in a subset of preparations that awake imaging is possible using ArcLight (**Figure S1**). Future experiments are needed to test the ability of ArcLight and other GEVIs to track higher-frequency respiratory dynamics in awake mice.

### GEVI comparison

We found that several newer GEVIs can report odor-evoked activity in the mouse olfactory bulb with a high signal-to-noise ratio when using epifluorescence imaging. Importantly, these sensors successfully captured the temporal heterogeneity previously observed across the rostral-caudal axis of the OB, where caudally located glomeruli exhibit stronger respiration-locking than those located rostrally (**Figure 7**) ^13–15^.

It is notable that while ArcLight had similar or lower signal-to-noise ratios than JEDI-2P and ASAP in some conditions, it consistently exhibited the brightest expression across each of the three conditions. The brightness of an *in vivo* sample is influenced by many experimental factors including the specific virus used, whether it is driven constitutively or in a Cre-dependent manner, and preparation variability. However, this result is anecdotally consistent with our previous experience with ArcLight as being relatively bright and stable ^13^. Because a sensor’s brightness dictates the total number of collected photons, it directly influences the achievable signal-to-noise ratio. A bright GEVI is particularly important for experiments involving 2-photon imaging, which has lower photon collection than what typically occurs under epifluorescence illumination ^25^. Therefore, future studies are needed to test whether the utility of other GEVIs under epifluorescence imaging conditions translates into successful *in vivo* 2-photon imaging ^3^.

### Conclusions

Our results demonstrate that 2-photon imaging of ArcLight expressed in MTC glomeruli has sufficient signal-to-noise ratio to image glomerular patterns of activation at high time resolution. Because GEVIs capture faster neuronal dynamics than GECIs, they are an ideal tool for experimental questions addressing precise temporal coding. For example, these sensors can be used to track the natural respiratory patterns present in awake mice, or to test current models of olfactory sensory processing which posit that the latency of the first olfactory response is the determining factor in achieving concentration-invariant odor perception ^26^.

## Author contributions

D.A.S. and L.L. were involved in the conception or design of the work. D.A.S. and L.L. were involved in the acquisition of the data for the work. L.L., E.L. and D.A.S. were involved in the analysis and interpretation of the results. L.L. and D.A.S. made the first drafts of all figures, and all authors were involved in revising them. D.A.S. wrote the first draft of the manuscript, and all authors were involved in revising it critically for important intellectual content. All authors approved the final version of the manuscript, agree to be accountable for all aspects of the work in ensuring that questions related to the accuracy or integrity of any part of the work are appropriately investigated and resolved. All persons designated as authors qualify for authorship, and all those who qualify for authorship are listed.

## Acknowledgements

NIDCD R01 DC020519 (D.A.S.), NSF CAREER # 2543400 (D.A.S.), NIDCD T32 DC000044 (E.L.), and funding through FSU’s Vice President for Research Chemosensory Postdoctoral Scholarship. Thanks to members of the Storace laboratory for helpful discussion on early drafts of this manuscript. All ASAP AAVs were gifts from the Michael Z. Lin lab at Stanford University.

## Data availability

All data are available to Neurophotonics and interested parties upon reasonable request.

---

**Figure S1:**
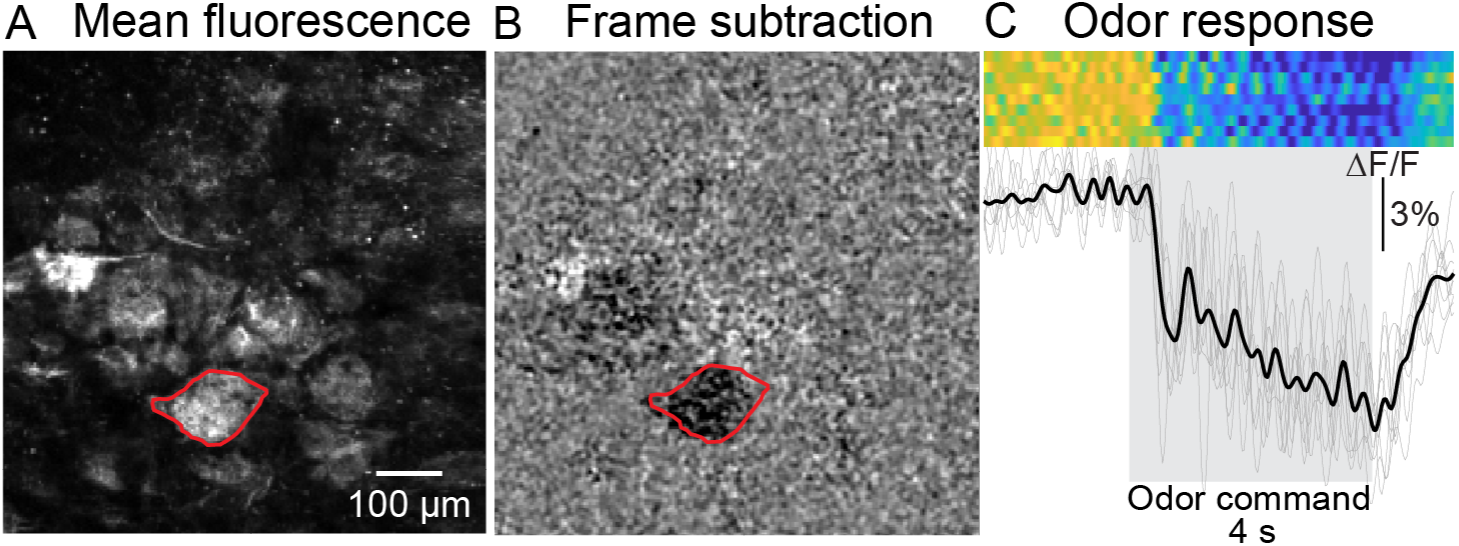
*In vivo* 2-photon awake imaging from ArcLight targeted to mitral/tufted glomeruli. (A-C) Mean fluorescence (A), frame subtraction, and fluorescence time course (C) from an exemplar preparation. The heat maps in panel C illustrate 9 single trials imaged from the region of interest illustrated in panels A-B. The response is to 6% methyl valerate.

## Bibliography

1. K. M. Igarashi et al., “Parallel mitral and tufted cell pathways route distinct odor information to different targets in the olfactory cortex,” J Neurosci 32(23), 7970–7985 (2012). 10.1523/jneurosci.0154-12.2012.

2. D. L. Sosulski et al., “Distinct representations of olfactory information in different cortical centres,” Nature 472(7342), 213–216 (2011). 10.1038/nature09868.

3. L. M. Leong and D. A. Storace, “Imaging different cell populations in the mouse olfactory bulb using the genetically encoded voltage indicator ArcLight,” Neurophotonics 11(3), 033402 (2024). 10.1117/1.NPh.11.3.033402.

4. D. A. Storace, L. B. Cohen and Y. Choi, “Using Genetically Encoded Voltage Indicators (GEVIs) to Study the Input-Output Transformation of the Mammalian Olfactory Bulb,” Front Cell Neurosci 13, 342 (2019). 10.3389/fncel.2019.00342.

5. D. Storace et al., “Toward Better Genetically Encoded Sensors of Membrane Potential,” Trends Neurosci 39(5), 277–289 (2016). 10.1016/j.tins.2016.02.005.

6. D. A. Storace and L. B. Cohen, “Measuring the olfactory bulb input-output transformation reveals a contribution to the perception of odorant concentration invariance,” Nat Commun 8(1), 81 (2017). 10.1038/s41467-017-00036-2.

7. J. Platisa et al., “Voltage imaging in the olfactory bulb using transgenic mouse lines expressing the genetically encoded voltage indicator ArcLight,” Sci Rep 12(1), 1875 (2022). 10.1038/s41598-021-04482-3.

8. L. M. Leong et al., “Evidence that interglomerular inhibition generates non-monotonic concentration-response relationships in mitral/tufted glomeruli in the mouse olfactory bulb,” J Physiol (2026). 10.1113/JP290366.

9. M. N. Economo, K. R. Hansen and M. Wachowiak, “Control of Mitral/Tufted Cell Output by Selective Inhibition among Olfactory Bulb Glomeruli,” Neuron 91(2), 397–411 (2016). 10.1016/j.neuron.2016.06.001.

10. A. K. Moran, T. P. Eiting and M. Wachowiak, “Circuit Contributions to Sensory-Driven Glutamatergic Drive of Olfactory Bulb Mitral and Tufted Cells During Odorant Inhalation,” Front Neural Circuits 15, 779056 (2021). 10.3389/fncir.2021.779056.

11. A. K. Moran, T. P. Eiting and M. Wachowiak, “Dynamics of Glutamatergic Drive Underlie Diverse Responses of Olfactory Bulb Outputs,” eNeuro 8(2)(2021). 10.1523/ENEURO.0110-21.2021.

12. N. Subramanian et al., “Recent odor experience selectively modulates olfactory sensitivity across the glomerular output in the mouse olfactory bulb,” Chem Senses 50(2025). 10.1093/chemse/bjae045.

13. D. A. Storace et al., “Monitoring brain activity with protein voltage and calcium sensors,” Sci Rep 5, 10212 (2015). 10.1038/srep10212.

14. M. Wachowiak et al., “Optical dissection of odor information processing in vivo using GCaMPs expressed in specified cell types of the olfactory bulb,” J Neurosci 33(12), 5285–5300 (2013). 10.1523/jneurosci.4824-12.2013.

15. H. Spors et al., “Temporal dynamics and latency patterns of receptor neuron input to the olfactory bulb,” J Neurosci 26(4), 1247–1259 (2006). https://doi.org/26/4/1247 [pii] 10.1523/JNEUROSCI.3100-05.2006.

16. T. P. Eiting and M. Wachowiak, “Differential Impacts of Repeated Sampling on Odor Representations by Genetically-Defined Mitral and Tufted Cell Subpopulations in the Mouse Olfactory Bulb,” J Neurosci 40(32), 6177–6188 (2020). 10.1523/jneurosci.0258-20.2020.

17. R. Shusterman et al., “Precise olfactory responses tile the sniff cycle,” Nat Neurosci 14(8), 1039–1044 (2011). 10.1038/nn.2877.

18. Y. B. Sirotin, R. Shusterman and D. Rinberg, “Neural Coding of Perceived Odor Intensity,” eNeuro 2(6)(2015). 10.1523/eneuro.0083-15.2015.

19. E. C. Sobel and D. W. Tank, “Timing of odor stimulation does not alter patterning of olfactory bulb unit activity in freely breathing rats,” J Neurophysiol 69(4), 1331–1337 (1993).

20. C. H. Yu et al., “The Cousa objective: a long-working distance air objective for multiphoton imaging in vivo,” Nat Methods 21(1), 132–141 (2024). 10.1038/s41592-023-02098-1.

21. M. Wachowiak et al., “Recalibrating Olfactory Neuroscience to the Range of Naturally Occurring Odor Concentrations,” J Neurosci 45(10)(2025). 10.1523/JNEUROSCI.1872-24.2024.

22. A. Dewan et al., “Single olfactory receptors set odor detection thresholds,” Nat Commun 9(1), 2887 (2018). 10.1038/s41467-018-05129-0.

23. L. Jennings et al., “The behavioral sensitivity of mice to acetate esters,” Chem Senses 47(2022). 10.1093/chemse/bjac017.

24. C. E. Johnson, E. Williams and A. Dewan, “Protocol for quantifying the odor detection threshold of mice,” STAR Protoc 4(4), 102635 (2023). 10.1016/j.xpro.2023.102635.

25. O. Braubach, L. B. Cohen and Y. Choi, “Historical Overview and General Methods of Membrane Potential Imaging,” Adv Exp Med Biol 859, 3–26 (2015). 10.1007/978-3-319-17641-3_1.

26. C. D. Wilson et al., “A primacy code for odor identity,” Nat Commun 8(1), 1477 (2017). 10.1038/s41467-017-01432-4.

